# Identification of Lipid Droplets in Gut Microbiota

**DOI:** 10.1101/2020.05.06.080317

**Authors:** Kai Zhang, Chang Zhou, Ziyun Zhou, Xuehan Li, Zemin Li, Mengwei Zhang, Xuelin Zhang, Congyan Zhang, Taotao Wei, Shuyan Zhang, Pingsheng Liu

## Abstract

Recent research has accumulating evidence to support that gut microbiota can be a newly discovered organ/tissue for humans. Although the functions of gut microbiota have been linked to almost all aspects of human physiological, most findings so far are still either belonged to correlational research or at preliminary stage of causal study due to the complexity of gut microbiota as well as lack of proper tools and methods. Thus, development of new approaches is essential and necessary. Lipid droplet is a cellular organelle governing cell lipid homeostasis that is directly related to human metabolic syndromes. Previously we and other groups found lipid droplets in several bacterial strains. We wonder if gut bacteria have lipid droplets. If so, these bacteria may play key roles to regulate gut lipid content, remove unnecessary lipids and produce useful lipid molecules for host. Here we identified lipid droplet-like structures in freshly isolated bacteria from mouse small and large intestines, as well as mouse and human feces. We also purified and analyzed lipid droplets from a cloned human gut bacterium *Streptomyces thermovulgaris*. Together, we demonstrate the existence of lipid droplets in gut microbiota and established an approach to study them.

## Results and Discussion

### LD-like structures in intestinal bacteria

Recent studies of gut microbiota have developed extremely fast and their composition and products have been associated with human physical and mental health[1–18]. Several sophistic approaches have been established to study gut microbiota, including high-throughput 16S rRNA sequencing, metagenomics, and massive bacterial cloning[19–27]. Since the number and species of gut microbiota are too huge, the resulting complexity is an obscure that makes their functions remain elusive[28]. More new techniques and approaches are required to dissect this important ecosystem[29, 30].

Lipid droplet (LD) as a cellular organelle has been found to play a key role in lipid homeostasis. The globular LD includes a neutral lipid core covered by a monolayer phospholipid membrane and some resident/dynamic proteins[31–33]. Except triacylglycerol (TAG), neutral lipids in LDs also include cholesterol ester, retinyl ester, wax ester, and polyhydroxyalkanoate (PHA)/polyhydroxybutyrate (PHB). Therefore, abnormal LD dynamics as well as ectopic lipid storage have been directly linked to many metabolic syndromes, such as obesity, fatty liver, and atherosclerosis, and diabetes[34–36]. We and others have previously reported that several strains of bacteria contain LDs and identified their main functions[37–40]. Consideration of how gut microbiota survives and what molecules they may produce in gut that affect human health, we decided to determine whether some gut bacteria have LDs. Thus, we first isolated bacteria freshly from mouse small and large intestine, and feces of mouse and human (Fig. 1A).

**Figure 1.**
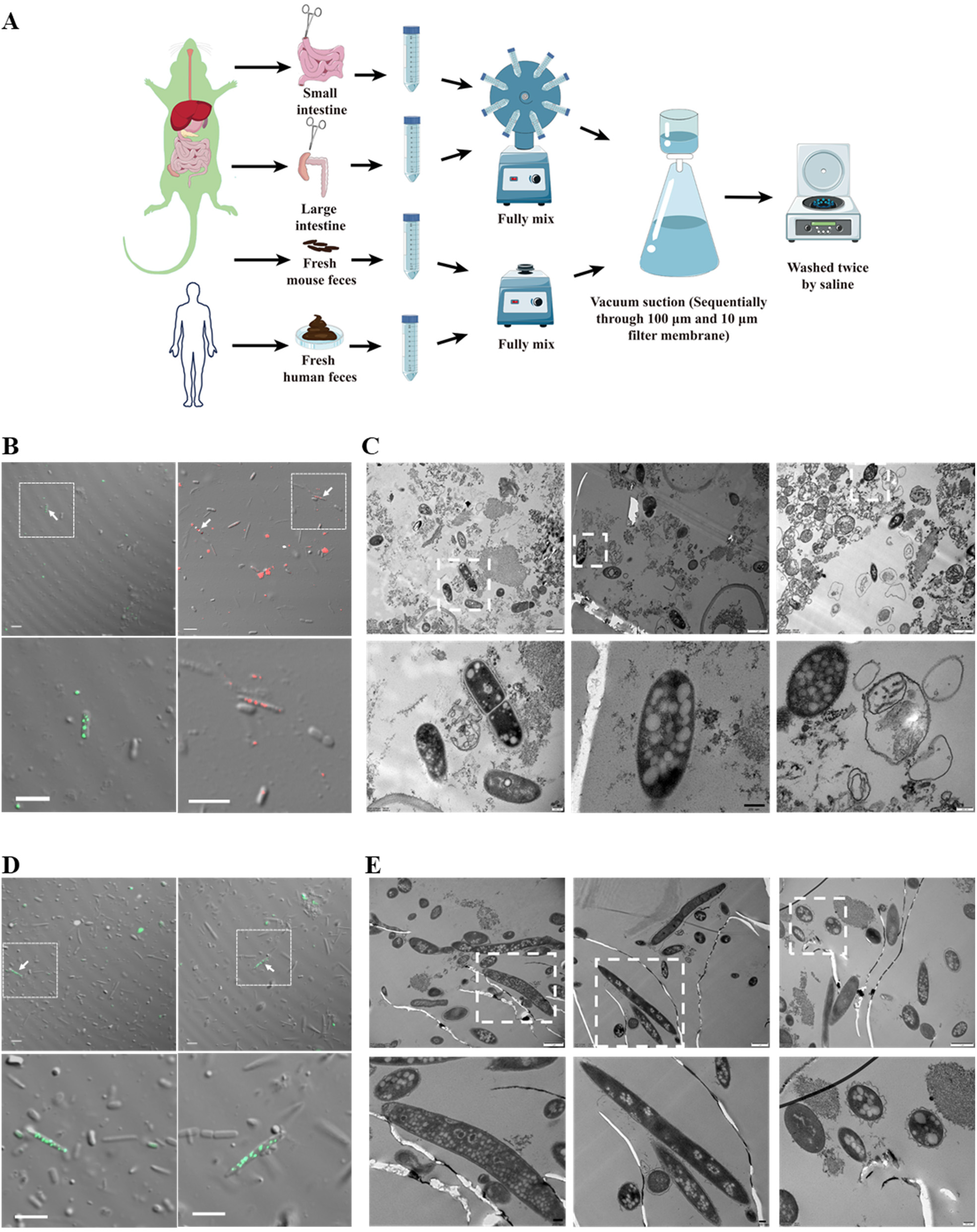
Lipid Droplet-like Structures in Mouse Small and Large Intestinal Microbiota. Gut microbiota was freshly isolated from mouse small and large intestines and subjected immediately to neutral lipid staining and sample preparation for transmission electron microscope (TEM). **A**. A workflow of fresh microbiota isolation from mouse small intestine, large intestine, mouse and human feces. In brief, mouse small and large intestine and their contents, and the mouse and human feces were collected respectively. Then, the samples were transferred into the 50 mL centrifuge tube and fully mixed in saline by vortex. Finally, the samples were filtered to remove food residue, and then the bacteria were concentrated and washed two times for subsequent experiments. Some figures were created using Servier Medical Art images (http://smart.servier.com). **B**. The freshly isolated microbiota from mouse small intestine were stained with LipidTOX Green or Red, and imaged by confocal microscope. Scale bar, 5 μm. **C**. Ultra-thin structure analysis of fresh microbiota from mouse small intestine by TEM. Briefly, the freshly isolated microbiota from mouse small intestine were fixed immediately and prepared into ultra-thin sections (70 nm) after dehydration in a series of ethanol, embedding into epoxy resin and sectioning. After staining, the sections were observed under TEM. Scale bar, 1 μm (upper panel) and 200 nm (enlarged, lower panel). **D**. The freshly isolated microbiota from mouse large intestine were stained with LipidTOX Green, and then imaged by confocal microscope. Scale bar, 5 μm. **E**. Ultrathin structure analysis of fresh microbiota from mouse large intestine by TEM. Briefly, the freshly isolated microbiota from mouse large intestine were fixed immediately and prepared into ultra-thin sections (70 nm) after dehydration in a series of ethanol, embedding into epoxy resin and sectioning. After staining, the sections were observed under TEM. Scale bar, 1 μm (upper panel) and 200 nm (enlarged, lower panel).

To determine existence of LD-like neutral lipid-containing structures, those freshly isolated bacteria were stained with a neutral lipid dye LipidTOX Red/Green immediately and observed under confocal microscope. Differential interference contrast (DIC) was applied to identify bacteria. Overlapping image represents neutral lipids inside of bacteria. After studies of three independent isolations, indeed, some mouse small intestinal bacteria showed positive fluorescent signals (Fig. 1B). Some fluorescent signals without overlapped bacterium-shaped structures in DIC were probably contaminations of staining (Fig. 1B, upper panel). Due to the resolution limit of confocal microscope, LipidTOX staining can only determine neutral lipids in bacteria without detailed structure information. To visualize morphology of these neutral lipid-containing structures, the freshly isolated bacteria were also fixed immediately and subjected to ultrastructural analysis using transmission electron microscopy (TEM). The TEM result revealed sphere-shaped structures with low density inside of some bacteria, verifying the existence of LD-like structures in those bacteria (Fig. 1C). In addition, both fluorescent signals and LD-like structures were found with different sizes and numbers, as well as in the bacteria that had different morphologies, suggesting the diversity of microbiota potentially containing LD in mouse small intestine.

The similar approach was applied on study of freshly isolated large intestinal microbiota. Figure 1D showed overlapping images of DIC and fluorescence signals. These findings were further verified by TEM analysis that revealed the sphere-shaped structures in the bacteria (Fig. 1E). These data present the neutral lipids and LD-like structures in freshly isolated bacteria from mouse small and large intestines, suggesting that LDs exist *in situ* in gut microbiota.

### LD-like structures in fecal bacteria

Most studies on gut microbiota so far have been carried out using fecal microbiota due to the difficulty in obtaining samples from human gut. We wonder if there are still some bacteria bearing LDs in feces, which can be important for study of small and large intestinal microbiota as well as for developing therapeutic treatment to remove neutral lipids such as TAG and cholesterol ester from gut. We, therefore, moved to determine if fecal microbiota also contain LDs. Bacteria from fresh feces of mouse and human were isolated using the method described (Fig. 1A). Similar to the studies on gut bacteria above, the freshly isolated fecal bacteria were subjected to identification of LD-like structures using both neutral lipid staining observed by confocal microscope and TEM analysis. Figure 2A showed that more bacteria were isolated from mouse feces compared to mouse intestine, and indeed, some of them contained multiple neutral lipid signals (LipidTOX Green). An aliquot of same bacterial isolation was analyzed by TEM and LD-like structures were clearly presented in some bacteria (Fig. 2B). As shown in Figure 2C, some LipidTOX Red fluorescent signals were overlapped with bacterial structures in DIC images in the freshly isolated bacteria from human feces. No surprise, the TEM also confirmed the existence of multiple LD-like structures in these bacteria (Fig. 2D). In agreement with gut microbiota shown in Figure 1, LD-like structures varied in size and number in fecal bacterial cells.

**Figure 2.**
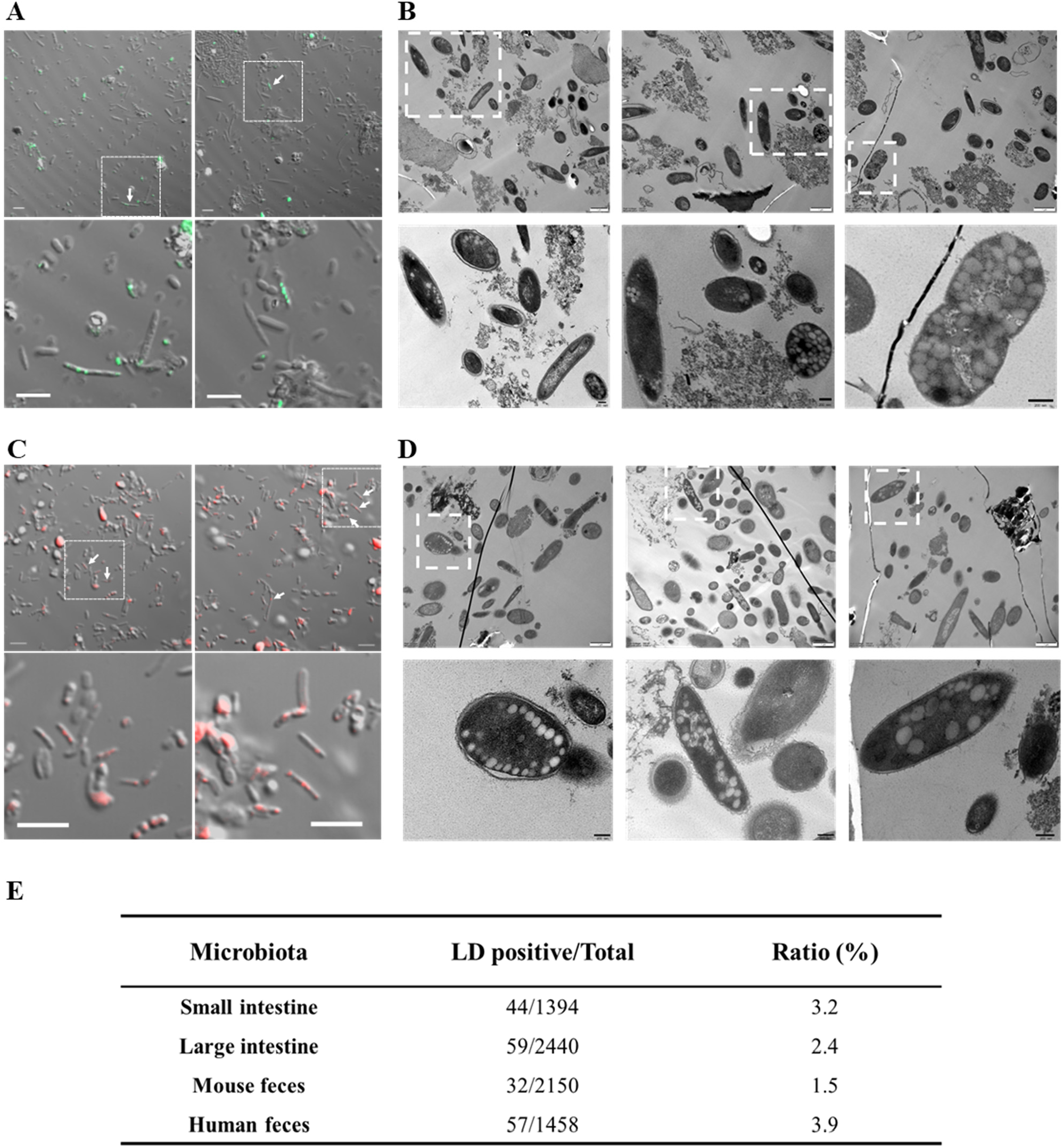
Lipid Droplet-like Structures in Mouse and Human Fecal Microbiota. Fecal microbiota was freshly isolated from mouse and human feces as previously described in Figure 1A and then subjected immediately to neutral lipid staining and sample preparation for TEM. **A**. The freshly isolated microbiota from mouse feces were immediately stained with LipidTOX Green, then mounted and imaged by confocal microscope. Scale bar, 5 μm. **B**. Ultra-thin structure analysis of fresh microbiota from mouse feces by TEM. Briefly, the microbiota from fresh mouse feces was fixed immediately after isolation and prepared into ultra-thin sections (70 nm) after dehydration in a series of ethanol, embedding into epoxy resin and sectioning. After staining, the sections were observed under TEM. Scale bar, 1 μm (upper panel) and 200 nm (enlarged, lower panel). **C**. The freshly isolated microbiota from human feces were stained with LipidTOX Red, and imaged by confocal microscope. Scale bar, 5 μm. **D**. Ultra-thin structure analysis of fresh microbiota from human feces by TEM. Briefly, the microbiota from fresh human feces was fixed immediately after isolation and prepared into ultra-thin sections (70 nm) after dehydration in a series of ethanol, embedding into epoxy resin and sectioning. After staining, the sections were observed under TEM. Scale bar, 1 μm (upper panel) and 200 nm (enlarged, lower panel). **E**. The ratio of Lipid droplet (LD)-positive microbiota from gut or feces. Each sample was counted from 20-50 confocal images. The ratio was calculated as the number of LD-positive bacteria to the number of total bacteria viewed.

Based on our collections, we counted 20-50 confocal images for each sample and calculated the ratio of LD-like structure positive bacteria as shown in Figure 2E. Together, even in a small population, LD-like structures could be detected in freshly isolated microbiota from small intestine, large intestine, and feces.

### Isolation of LDs from a gut bacterium

To demonstrate that these LD-like structures are LDs, 5 cloned human gut bacteria were collected, cultivated, and analyzed. One of them, *Streptomyces thermovulgaris* was cultivated to stationary phase (Fig. 3Aa) and subjected to TAG measurement (Fig. 3Ab). Their content of TAG was measured using TAG assay kit and showed in Figure 3Ab, 68 μg/mg protein. Then the LD-like structures that accumulate neutral lipids were studied using LipidTOX staining plus confocal imaging and ultrastructural analysis using TEM (Fig. 3B and C). *Streptomyces thermovulgaris* was previously isolated from human fecal samples[41, 42] and can be probiotics since its thermostable endo-xylanase is able to produce xylooligosaccharides (XOs) from corncob[43]. XOs from fruits and vegetables are beneficial to animal health through enhancing the population of bacteria that produce short chain fatty acids[44]. These previous studies indicate that *Streptomyces thermovulgaris* is an important and useful strain of human microbiota. Therefore, LDs were isolated from the bacterium using our established method[37]. The isolated LDs were verified by neutral lipid staining and size distribution analysis (Fig. 3Da and b). Their lipids were then analyzed using thin layer chromatography (TLC) and three major neutral lipids were identified, including TAG, unknown lipids, and PHB (Fig. 3Dc, lane 6). Before being subjected to MS analysis, reproducibility of LD isolation was determined. Based on comparison of two isolations, LD protein compositions were almost identical (Fig. 4Aa, lanes 1 and 2). In addition, LD protein composition was unique compared to other cell fractions such as cytosol (Cytosol), whole cell lysate (WCL), and total membrane (TM) (Fig. 4Aa). Together, these experiments confirmed that a human gut bacterium *Streptomyces thermovulgaris* had LDs, supporting above findings in mouse and human microbiota (Figs. 1 and 2).

**Figure 3.**
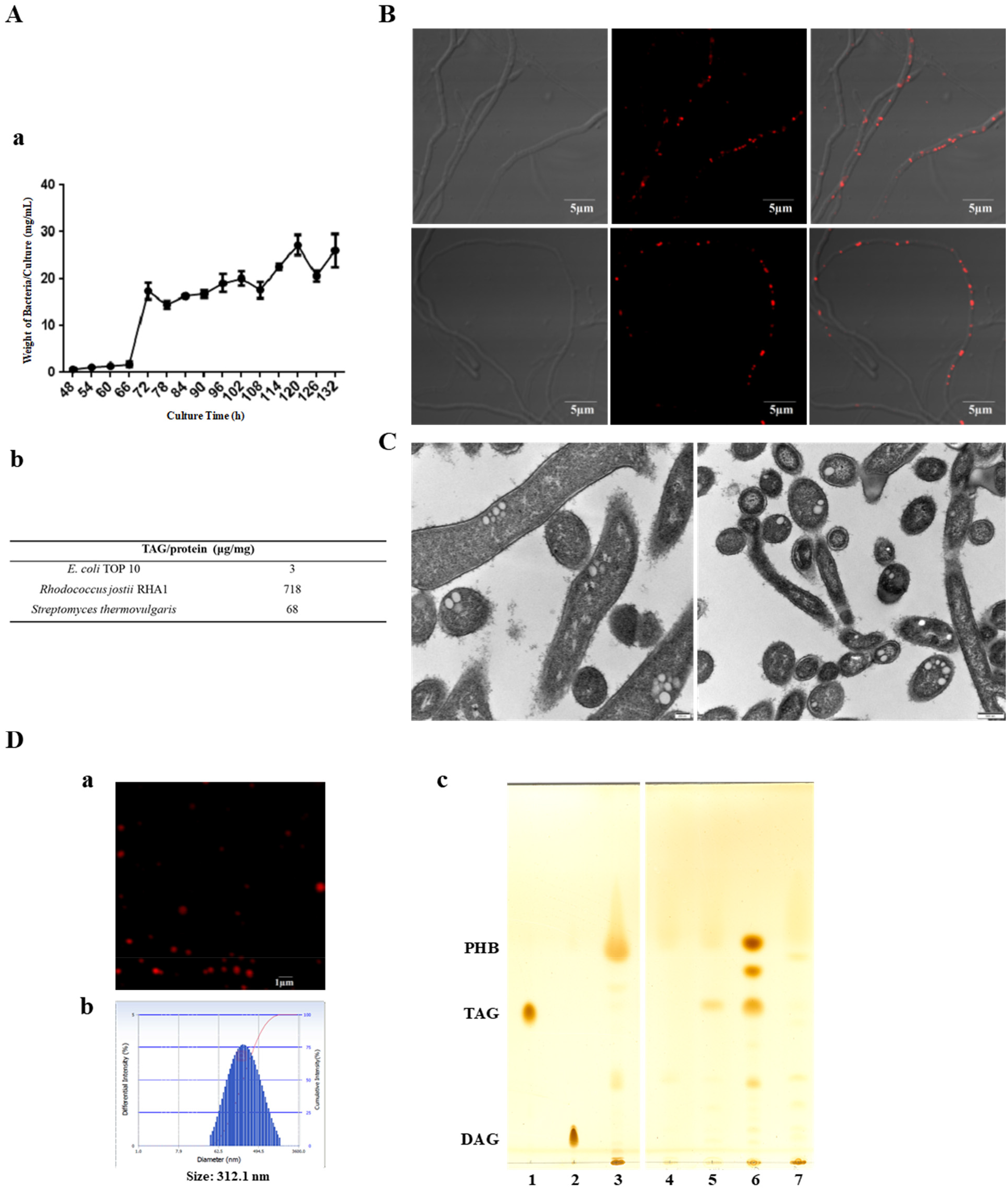
Identification of Lipid Droplet in Cloned Gut Bacterium *Streptomyces thermovulgaris*. Cloned human gut bacterium *Streptomyces thermovulgaris* was cultured in ISP2 medium. **A**. The growth curve of *Streptomyces thermovulgaris* cultured in ISP2 medium at 37°C was monitored using dry weight method every 6 h (**a**). Relative TAG content was measured by an enzymatic assay in *Streptomyces thermovulgaris* when it grew to 120 h (**b**). **B**. Confocal images of *Streptomyces thermovulgaris* were captured after neutral lipid staining with LipidTOX Red. Scale bar, 5 μm. **C**. Ultra-thin structure analysis of *Streptomyces thermovulgaris* by TEM. Briefly, the bacteria were fixed, dehydrated, embedded immediately after collection, and finally sectioned into ultrathin sections (70 nm). The sections were stained and viewed in the TEM. Scale bar, 200 nm (left panel) and 500 nm (right panel). **D**. Lipid droplets (LDs) were isolated from *Streptomyces thermovulgaris* using the method that we previously reported. Isolated LDs were stained with LipidTOX Red and observed by confocal imaging (**a**). The diameter of isolated LDs was analyzed by dynamic light scattering (**b**). Neutral lipids of isolated LDs were analyzed by TLC. Lane 1, TAG; Lane 2, diacylglycerol (DAG); Lane 3, PHB; Lane 4, whole cell lysate of *Escherichia coli*; Lane 5, whole cell lysate of *Rhodococcus jostii*; Lane 6, LDs of *Streptomyces thermovulgaris*; Lane 7. whole cell lysate of *Streptomyces thermovulgaris* (**c**).

**Figure 4.**
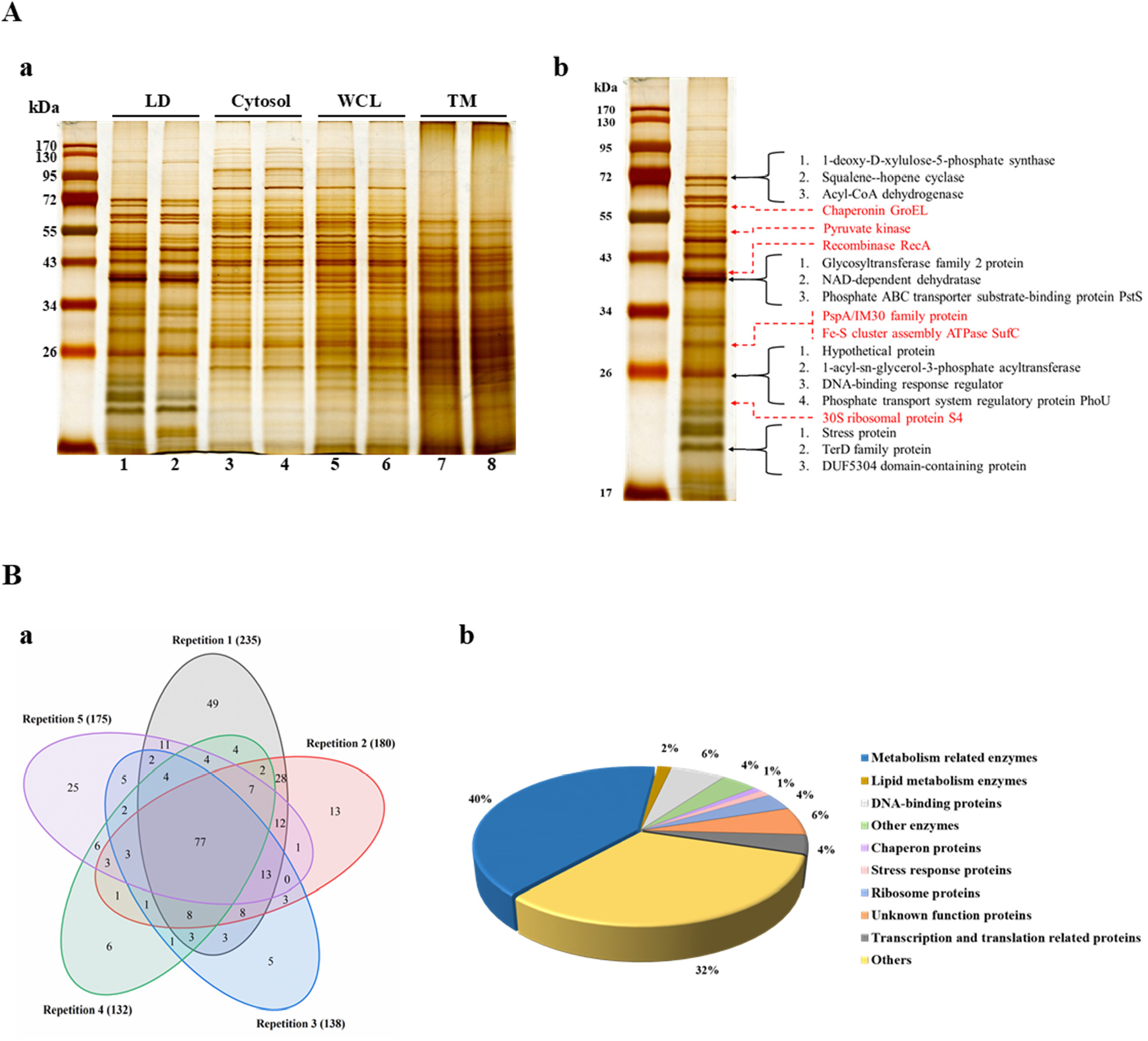
Proteomic Analysis of Isolated Lipid Droplets from Cloned Gut Bacterium *Streptomyces thermovulgaris*. *Streptomyces thermovulgaris* was cultured in ISP2 medium for 120 h. Lipid droplets (LDs) were isolated according to the method established previously. Briefly, the bacteria were collected and homogenized through a French pressure cell at 1,200 bar. LDs were separated by sucrose density gradient ultracentrifugation. **A**. The protein profiles of the cellular fractions were analyzed by SDS-PAGE and silver staining. LD, lipid droplet; WCL, whole cell lysate; TM, total membrane. **B**. Four major bands in the gel as arrow indicated were sliced and subjected to in-gel digestion followed by LC-MS/MS identification. Besides, the total LD proteins were also subjected to proteomic analysis directly and several abundant proteins identified were indicated using dotted lines. **C**. Five LD isolations from five independent cultures were conducted respectively. The total LD proteins were subjected to proteomic identification as five biological replicates. A Venn diagram was built to present the overlap of LD proteomes from the five biological replicates (**a**). The identified LD proteins from LC-MS/MS were then analyzed using bioinformatics tools and the proteins with 2 unique peptides or above were selected for further study. Based on their functions, they were categorized into 10 groups (**b**).

### Proteomic analysis of isolated LDs

To determine major proteins of isolated LDs from human microbiota, the four major bands indicated in Figure 4Ab (adapted from Fig. 4Aa, lane 1) were sliced and subjected to LC/MS/MS identification. The identified major proteins from four bands were listed in the right side of the PAGE (in black). Then the proteins of isolated LDs were subjected to LC/MS/MS analysis using shotgun method. For the accuracy of *Streptomyces thermovulgaris* LD proteome, five replicates had been carried out including five cultures, five LD isolations, and five proteomic identifications. The number of proteins existed in all five proteomes was 77 (Fig. 4Ba). The number ofidentified proteins with two unique peptides was 310 and these proteins were categorized into 10 groups (Fig. 4Ba). Phage shock protein A (PspA), a previously identified LD major protein of bacteria[37, 40], and other major proteins were also listed in Figure 4Ab (in red).

We demonstrated the existence of LDs in mouse and human gut microbiota, and provided a proteome of LD from human gut microbiota. This new method/approach, combined with other LD research work, allows us to investigate gut microbiota on a completely new angle. Based on current understandings of the organelle, LDs in human cells have significant impact on human health. It can be expected that LDs in gut microbiota also play key roles in host gut functions including food digestion and absorption, essential molecule production, gut environment, immune protection, and fecal energy output, which is also fundamental for human health.

## Materials and Methods

### Materials and bacterial strain

The LipidTOX Red/Green were from Invitrogen. Sodium oleate and lipid strandards were from Sigma. 25% glutaraldehyde solution (EM grade), 8% paraformaldehyde solution (EM grade), Embed 812 kit, uranyl acetate and lead citrate were all purchased from Electron Microscopy Sciences (Hatfield, USA). The strain *Streptomyces thermovulgaris* was purchased from Institute of Microbiology, Chinese Academy of Sciences.

### Freshly isolation of microbiota from mouse gut and feces

All animal experiments were approved by the Committee of Biosafety, Ethics and Experimental Animal Management of Institute of Biophysics, Chinese Academy of Sciences, permit number SYXK (Jing) 2016-0026, and performed in accordance with the NIH Guide for the Care and Use of Laboratory Animals (8th edition). Eight-week-old C57BL/6 mice were purchased from Beijing Vital River Laboratories. The mice were given ad libitum access to chow diet and water.

Two mice were used in each experimental group. The gut microbiota was isolated according to the method previously reported with modification[45]. Briefly, the fresh feces (1 fecal sample: minimum 0.2 g) were collected 10 min before decapitation. After decapitation, small intestine and large intestine were entirely taken out for use, and the mesentery was removed with careful dissection. The small intestine and large intestine were carefully slit open with surgical scissors. The contents of small intestine and large intestine, as well as mouse feces were transferred immediately into saline (4°C, 20 mL) in 50 mL centrifuge tube respectively and rotated for 10 min. The corresponding mixture was filtered through a suction filtration system with a 100-μm membrane and subsequently a 10-μm membrane filter. The resulting sample solution was centrifuged at 3,200*g* for 20 min at 4°C. The pellet was resuspended with 1 mL cold saline and transferred into 1.5 mL centrifuge tube and centrifuged at 10,000*g* for 2 min at 4°C. The washing step was repeated once more. The resulting bacteria were for further analysis.

### Isolation of bacteria from fresh human feces

Bacteria of fresh human feces were isolated similarly to the method described above. Fresh fecal sample (about 10 mL) was collected from a 26-year-old healthy man. The volunteer had no previous history of gastrointestinal disease and had not consumed antibiotics in the three months prior to this study. The fresh feces were transferred into 50 mL centrifuge tube immediately and resuspended with 30 mL cold saline. The sample was vigorously vortexed 5 times for 1 min, with 1-min intervals in between vortexing. The mixture was passed through a suction filtration system with a 100-μm membrane and subsequently a 10-μm membrane filter. The bacteria were collected and washed similarly to the procedure described above for mouse samples.

### LipidTOX staining of isolated bacteria

Freshly isolated microbiota was added onto cover glasses pretreated with poly-L-lysine (PB0589) before washing. The microbiota was then stained with LipidTOX (1:500, v/v) for 30 min. The cover glasses were then mounted on glass slides using mounting media (P0126) and observed with a ZEISS LSM 980 confocal laser scanning microscope.

### Ultrastructural analysis of bacteria by transmission electron microscopy (TEM)

Ultrastructure of bacteria, both freshly isolated and cultured bacteria, were analyzed using TEM after ultra-thin sectioning. Briefly, bacteria were pelleted and washed twice with indicated buffer. For freshly isolated bacteria from gut and feces, 0.1 M PB (pH 7.2) was used and for *in vitro* cultured bacteria, 50 mM K-Pi (pH 7.2) was used. Then the cells were fixed in buffered glutaraldehyde (2.5%, v/v) and paraformaldehyde (2%, v/v) overnight at 4°C. Subsequently, the cells were further fixed in 2% (w/v) potassium permanganate for 5 min at room temperature. Then the samples were dehydrated in an ascending concentration series of ethanol followed by propylene oxide at room temperature. After dehydration, the samples were embedded in Embed 812 and prepared as 70-nm-thick ultrathin sections using Leica EM UC6 Ultramicrotome. Ultrathin sections were collected on formvar-coated copper grids and stained with uranyl acetate and lead citrate. The sections were then observed with Tecnai Spirit electron microscope (FEI, Netherlands).

### Cultivation and growth curve of *Streptomyces thermovulgaris*

*Streptomyces thermovulgaris* was cultured in 10 mL ISP2 medium (1% Malt extract, 0.4% Yeast extract, 0.4% D-glucose, pH 7.2) at 37°C. The growth of *Streptomyces thermovulgaris* was monitored by determination of its dry weight. At each time point, three parallel cultures were recorded.

### Lipid droplet purification

Lipid droplets of *Streptomyces thermovulgaris* were purified according to our previously established method with modification[37]. In brief, bacteria were collected and washed twice with Buffer A (25 mM tricine, 250 mM sucrose, pH 7.8). After incubation in Buffer A for 20 min on ice, the bacteria were homogenized by passing through a French pressure cell six times at 1,200 bar, 4°C. The sample was then centrifuged at 6,000*g* for 10 min to remove cell debris and unbroken cells. The resulting supernatant (10 mL) was overlaid with 2 mL Buffer B (20 mM HEPES, 100 mM KCl, 2mM MgCl_2_, pH 7.4) and centrifuged at 182,000*g* for 1.5 h at 4°C (Beckman SW40). Then the lipid droplets on the top were collected and washed for further analysis.

### Analysis of size and staining of isolated LDs

The size of LDs was analyzed by a BECKMAN Delsa Nano C particle analyzer (Thermo, USA). Purified LDs were stained with LipidTOX (1:500, v/v) for 30 min and mounted on glass slide for observation with Olympus FV1200 Imaging System.

### Analysis of neutral lipids by thin layer chromatography (TLC)

After collection, LDs were treated with 1 mL of chloroform: acetone (8:2, v/v) and vigorously vortexed. Then the tube was centrifuged at 20,000*g* for 10 min at 4°C. The protein pellet was dissolved in 2 × SDS sample buffer for further analysis. The supernatant was transferred to a new tube and evaporated under a stream of nitrogen. The resulting lipid was dissolved in chloroform for TLC analysis.

Collected bacteria were resuspended with 400 μL PBS. Then 400 μL methanol and 800 μL chloroform were added. The sample was vigorously vortexed 3 times for 1 min, with 1-minute intervals in between vortexing. Then the sample was centrifuged at 20,000*g* for 10 min at 4°C and the lower phase was transferred into a new tube. The remaining sample was extracted with 800 μL chloroform again. The two lipid solutions were combined and evaporated under a stream of nitrogen. The resulting lipid was dissolved in chloroform for TLC analysis.

The lipids were loaded on TLC plate and developed using solvent of hexane-diethyl ether-acetic acid (70:30:1, v/v/v). Then the TLC plate was stained with iodine vapor.

### Triacylglycerol concentration assay

The content of bacterial TAG was measured by using a Triglycerides Kit (GPO-PAP Method), according to the manufacturer’s protocol (Biosino Bio-Technology and Science, China). The BCA kit (Thermo, USA) was used for measurement of protein content.

### Silver staining

The gel was fixed for 30 min and transferred into sensitization solution for another 30 min. Following this step, the gel was washed four times, 5 min each time then incubated with silver nitrate solution for 20 min. The gel was rinsed briefly with double distilled water followed by incubation in 2.5% (w/v) anhydrous sodium carbonate solution until protein bands are conspicuously satisfactory. The reaction was stopped with disodium ethylenediaminetetraacetic acid solution.

### Mass spectrometry study

For extracted proteins from isolated LDs, the proteins were dissolved in 8 M urea. For the protein bands indicated in Figure 4Ab, the bands were excised from gel, cut into small plugs, and washed twice with water. The gel pieces were destained with 40% acetonitrile/50 mM NH_4_HCO_3_, and then dehydrated with 100% acetonitrile and dried for 5 min using a Speedvac.

After reduction with dithiothreitol and alkylation with iodoacetamide, the samples were digested with trypsin at 37°C overnight. After quenching the reaction with formic acid, the peptides were desalted, followed by vacuum centrifugation. The resulting peptide mixtures were then transferred for LC-MS/MS analysis.

All analyses were performed on nanoLC-LTQ-Orbitrap XL mass spectrometer (Thermo, San Jose, CA). For nanoLC, Easy n-LC 1200 system was equipped with 30 mm ReproSil-Pur C18-AQ (Dr. Maisch GmbH, Ammerbuch, Germany) trapping column (packed in-house, i.d.150 μm; resin, 5 μm) and 150 mm ReproSil-Pur C18-AQ (Dr. Maisch GmbH, Ammerbuch, Germany) analytical column (packed in-house, i.d. 75 μm; resin, 3 μm). Solvents used were 0.5% formic acid water solution (buffer A) and 0.5 % formic acid acetonitrile solution (buffer B). Elution was achieved with a gradient of 5–10% B in 3 min, 10–28% B in 70 min, 28–40% B in 8 min, 40–100% B in 2 min, 100% B for 7 min, at a flow rate of 300 nL/min.

Eluting peptide cations were converted to gas-phase ions by Nanospray Flex ion source at 2.1 kV. The heated capillary temperature was 225°C. The mass spectrometer was operated in a data-dependent mode to switch automatically between MS and MS/MS. Survey full scan MS spectra were acquired from *m/z* 300 to *m/z* 1800, and the 10 most intense ions with charge state above 2 and above an intensity threshold 500 were fragmented in the linear ion trap using normalized collision energy of 35%. For the Orbitrap, the AGC target value was set 1e6 and a maximum fill time for full MS was set 500 ms. Fragment ion spectra were acquired in the LTQ with AGC target value of 3e4 and a maximum fill time of 150 ms. Dynamic exclusion for selected precursor ions was set at 90 s. Lock mass option was enabled for the 445.120025 ion.

The raw data was processed using Proteome Discoverer (version 1.4.0.288, Thermo Fischer Scientific). MS2 spectra were searched with SEQUEST engine against *Streptomyces thermovulgaris* database and contaminant protein database. Database searches were performed with the following parameters: precursor mass tolerance 20 ppm; MS/MS mass tolerance 0.6 Da; two missed cleavages for tryptic peptides; methionine oxidation as variable modification; cysteine carbamidomethylation as fixed modification. The resulting results were filtered for a 1% false discovery rate (FDR) at the PSM level utilizing the percolator-based algorithm. Peptide identifications were grouped into proteins according to the law of parsimony.

The raw data of four bands were subjected to proteomic analysis, and the major proteins were screened based on the Molecular Weight, PSMs and Unique Peptides. The data from five biological replicates of shotgun proteomes were screened for proteins with two unique peptides or above, respectively. Then the proteins existed in all five proteomes were screened. Also the identified proteins from the five replicates were combined, manually assigned based on NCBI database (https://www.ncbi.nlm.nih.gov/) and UniProt (https://www.uniprot.org/) to search the functional categorization of those LD-associated proteins.

## Acknowledgments

The authors thank Ms. Yan Teng for her help of taking confocal images and Ms. Zhensheng Xie for her technical supports of proteomic study. This work was supported by the National Key R&D Program of China (Grant No. 2016YFA0500100, 2018YFA0800900 and 2018YFA0800700), National Natural Science Foundation of China (Grant No. 91857201, 31671402, 91954108, 31671233, 31771314, 31701018 and U1702288). This work was also supported by the “Personalized Medicines——Molecular Signature-based Drug Discovery and Development”, Strategic Priority Research Program of the Chinese Academy of Sciences, Grant No. XDA12040218.

## Author contributions

P.L. and S.Z. designed the project. K.Z., S.Z., C.Z., and X.L. performed the experiments. Z.Z. carried out bioinformatic analysis. Z.L. and M.Z. helped with triacylglycerol assay. X.Z., C.Z. and T.W. helped with MS study. P.L., S.Z., and C.Z. wrote the manuscript. All authors have read and approved the manuscript.

